# Reinvestigation of grain weight genes *TaTGW6* and *OsTGW6* casts doubt on their role in auxin regulation in developing grains

**DOI:** 10.1101/2020.10.14.340042

**Authors:** Muhammed Rezwan Kabir, Heather M. Nonhebel

**Affiliations:** School of Science and Technology, University of New England, Armidale, New South Wales 2351, Australia

**Keywords:** Auxin, grain weight, IAA-Glc, OsSTRL2, *TGW6*, wheat

## Abstract

The *THOUSAND-GRAIN WEIGHT 6* genes (*TaTGW6* and *OsTGW6*) are reported to result in larger grains of wheat and rice by reducing production of indole-3-acetic acid (IAA) in developing grains. However, a critical comparison of data on *TaTGW6* and *OsTGW6* with other reports on IAA synthesis in cereal grains requires that this hypothesis be reinvestigated. Here, we show that *TaTGW6* and *OsTGW6* are members of a large gene family that has undergone major, lineage-specific gene expansion. Wheat has nine genes, and rice three genes encoding proteins with more than 80% amino acid identity with TGW6 making it difficult to envisage how a single inactive allele could have a major effect on IAA levels. TGW6 is proposed to affect auxin levels by catalysing the hydrolysis of IAA-glucose (IAA-Glc). However, we show that developing wheat grains contain undetectable levels of ester IAA in comparison to free IAA and do not express an IAA-glucose synthase. Previous work on *TGW6*, reported maximal expression at 20 days after anthesis (DAA) in wheat and 2 DAA in rice. However, we show that neither gene is expressed in developing grains. Instead, *TaTGW6, OsTGW6* and their close homologues are exclusively expressed in pre-emergence inflorescences; *TaTGW6* is expressed particularly in microspores prior to mitosis. This combined with evidence for high levels of IAA production from tryptophan in developing grains demonstrates *TaTGW6* and *OsTGW6* cannot regulate grain size via the hydrolysis of IAA-Glc. Instead, their similarity to rice strictosidine synthase-like (*OsSTRL2*) suggests they play a key role in pollen development.

## Introduction

Grain size is an important component of cereal yield and grain quality that has been the subject of extensive research in all cereals including wheat (Beral et al. 2020). This work has identified a large number of quantitative trait loci (QTLs) for grain weight in wheat (Kumar et al. 2019; Sukumaran et al. 2018), although most have not been fully validated and cloned. Nevertheless a number of candidate genes for grain size and grain weight have been reported and listed in reviews such as Cao et al. (2020), Gupta et al. (2020) and Li and Li (2016). Brinton and Uauy (2019) have argued the importance of rigorously confirming the mechanistic role of candidate genes within QTLs to establish they are directly responsible for influencing grain weight as well as to inform their interaction with other factors. A number of grain size and grain weight genes are associated with plant hormones including *TaCKX6-D1, TaTGW6-A1* and *TaTGW-7A* (Gupta et al. 2020). However, although plant hormones have been argued to play a major role in early grain development, the mechanism of their involvement is still largely unknown (Basunia and Nonhebel 2019). In particular, there are contradictory reports relating to the role of indole-3-acetic acid (IAA) in cereal grain development. Both, *TaTGW6* and *TaTGW-7A* are inactive alleles, reported to increase grain size by reducing the level of auxin in developing grains (Hu et al. 2016a; Hu et al. 2016b). This claim requires further investigation as other work suggests IAA production has a positive correlation with grain fill in wheat (Li et al. 2014; Shao et al. 2017). In other cereals, *Big grain1* increases rice grain size via its effect on auxin transport that results in increased IAA in the panicles (Liu et al. 2015). In addition, the rice *tillering and small grain 1* (*tsg1*) mutant with reduced IAA biosynthesis has small grains (Guo et al. 2019) and the *defective endosperm18* (*de18*) mutant of maize is specifically deficient in endosperm IAA production (Bernardi et al. 2012).

*TGW6* was first identified in rice following high resolution mapping of a major QTL for thousand-grain weight (TGW) using backcrossed inbred lines produced from Nipponbare and the Indian landrace Kasalath (Ishimaru et al. 2013). The authors noted a single open reading frame within the mapped location which had a single base pair deletion compared to the Nipponbare allele. Transformation of an RNAi construct for *TGW6* into Nipponbare and NIL(*TGW6*) confirmed that *TGW6* was responsible for the increased grain size. Expression analysis indicated that the gene was active in leaf tissue as well as the panicles, with maximum upregulation reported at 2 days after anthesis (DAA). The authors suggested the allele might increase grain length via effects on the timing of endosperm cellularisation and noted that auxin may affect the timing of endosperm cellularisation. This combined with analysis of the amino acid sequence of TGW6 and molecular modelling studies led the authors to suggest that it might have IAA-glucose (IAA-Glc) hydrolase activity. This was confirmed using cloned Nipponbare TGW6. Supporting this result, the IAA content of grains at 3 DAA was reduced in the NIL(*TGW6*) compared to Nipponbare and auxin treatment of ovaries at the start of flowering reduced the endosperm length at 5 DAA. Ishimaru et al. (2013) thus appear to have built a strong case for the proposed role of TGW6. Later an orthologous gene *TaTGW6* was reported in wheat by Hu et al. (2016a). As was the case with rice, the null allele, *TaTGW6-c*, as well as a mutant allele, *TaTGW6-b* were associated with higher grain weight and lower IAA content of the grains. Nevertheless a closer look at both papers as well as consideration of major differences between them raises a number of key questions that need to be addressed.

Firstly, in order to have the reported effect on the IAA content of grains, the major source of IAA would have to be hydrolysis of the IAA-Glc conjugate rather than *de novo* synthesis from tryptophan via tryptophan aminotransferase (TAR) and indole-3-pyruvate monooxygenase (YUCCA). Neither Ishimaru et al. (2013) nor Hu et al. (2016a) investigated the availability of IAA-Glc as a substrate for TGW6. Furthermore, the TAR/YUCCA pathway of IAA production is strongly upregulated in rice from 4 DAA (Abu-Zaitoon et al. 2012) and from 10 DAA in wheat (Kabir et al. 2020). The *TaTGW6* work has further problems in that the authors did not assess the activity of the gene product and the IAA measurements used inappropriate, low specificity methods (High performance liquid chromatography with UV absorbance detector). Hu et al. (2016a) also reported that *TaTGW6* was expressed much later in grain development, close to the end of the grain fill period at 20 DAA. The differences in IAA content of grains were similarly detected late in grain development. Thus the wheat gene would have to affect grain size by a completely different mechanism. Finally, although Hu et al. (2016a) identified a large number of genes with high homology to *OsTGW6* in a combination of *Triticum Urartu* (source of wheat A genome)*, Aegilops tauschii* (source of wheat D genome) and *Hordeum vulgare* (barley), they reported that in hexaploid wheat, only *TaTGW6* was associated with grain weight. It is unclear how a single inactive allele from a group of highly homologous genes could have a major effect on IAA content via hydrolysis of IAA-Glc.

We therefore set out to explore two major questions raised above in relation to *TaTGW6*. 1. How many *TaTGW6-*like genes are expressed in developing wheat grains? 2. Is IAA-Glc present in developing wheat grains? We also investigated whether rice also has multiple *TGW6-*like genes. Finally, we set out to reinvestigate the timing and location of expression of *TGW6-*like genes in rice and wheat.

## Materials and methods

### Bioinformatic analysis

Wheat (*Triticum aestivum*) protein sequences homologous to OsTGW6, OsIAGLU, ZmIAGLU query sequences were downloaded from IWGSC RefSeq v1.1. via EnsemblPlants 47 (Kersey et al. 2015) following BlastP searches. Rice TGW6-like protein sequences were downloaded from Phytozome 12 (Goodstein et al. 2012). Phylogenetic analysis of protein sequences was carried out in MEGA7.0.26 (Kumar et al. 2016) using the Maximum Likelihood method (Jones et al. 1992). Multiple sequence alignment (MSA) were performed using MUSCLE (Edgar 2004). Bootstrap confidence levels were obtained using 500 replicates (Felsenstein 1985). Evolutionary distances were computed using Poisson correction method (Zuckerkandl and Pauling 1965). Pairwise sequence alignment to determine amino acid identity was performed by EMBOSS Needle (Emery and Morgan 2017). Global expression of *TaTGW6-*like genes and putative *TaIAGLU* genes was investigated using RNA sequencing (RNA-seq) data available in expVIP (http://www.wheat-expression.com). Rice RNA-seq data were obtained from Rice Genome Annotation Project (Kawahara et al. 2013).

### Plant materials

Plants (Chinese Spring wheat variety *Triticum aestivum* L.) were grown in 20 cm × 20 cm pots under natural light at 23/14°C (day/night) in the glasshouse at the University of New England, Armidale, NSW. Pots were fertilized once a week with Thrive^®^ (Yates, 1 g/L) from the initiation of tillering stage. Spikes were tagged when the first spikelet reached anthesis. Samples were collected from nine-day-old whole seedlings, pre-anthesis stages (40-50 mg), and at 5, 10, 15, 20 and 30 DAA from grains (70-90 mg). All samples were harvested at the same time each day (4:00 to 5:00 pm), then snap-frozen in liquid nitrogen and stored at −80°C. All analyses were carried out on at least three independent biological replicates harvested from different plants, on different days.

### RNA extraction and quantitative analysis

Wheat grain samples were ground in liquid nitrogen and total RNA was extracted using Trizol (Invitrogen). RNA concentration and 260/280 ratio were determined using a NanoDrop ND-8000 Spectrophotometer (Thermo Scientific). RNA samples with A_260/280_ of 1.8-2.0 were used for further analysis. The RNA quality was checked by agarose gel electrophoresis for two clear bands of 18S and 28S rRNAs (Nolan et al. 2006). Due to the large number of genes from each genome, 13 primer sets for *TaTGW6* and two for *TaIAGLU* were designed to amplify groups of highly similar genes as shown in Table S1. All melting points were in the range of 58–60°C and product sizes between 100–250 bp. Amplification was carried out using a One-Step RT-PCR Kit (QIAGEN) and BIO-RAD T100 Thermal Cycler, with gel analysis to confirm a single product of the expected size. Controls with no reverse transcriptase and no template were included with each set of reactions. In addition, positive controls confirmed that primer sets were able to amplify DNA samples.

### Ester and free IAA measurement

Wheat grain samples (70–90 mg) were ground in liquid nitrogen; 200 μL of 65% isopropanol /35% 0.2 M pH 7.0 imidazole buffer (Chen et al. 1988) was then added with [^13^C_6_] IAA internal standard (Cambridge Isotope Laboratories Inc.), and samples were extracted on ice for 1 h. Amounts of standard added varied with the age of samples; 16 ng of [^13^C_6_] IAA was added to pre-anthesis samples and 78 ng of [^13^C_6_] IAA was added to 20 and 30 DAA samples. Blank samples without plant tissue were taken through the entire extraction and analysis protocol to ensure that no contamination from the unlabelled IAA in the laboratory occurred.

Following extraction, samples were diluted with 1 mL deionized water and divided in two separate tubes. One portion from each extract was processed for ester IAA, the other for free IAA. An additional 0.5 mL deionized water was added to each tube, which was then centrifuged and the supernatant was transferred to fresh tubes. Ester IAA samples were transferred to 0.5 mL centrifugal filters, with 3 kDa molecular weight cut off (Amicon^®^ Ultra, Merck Millipore Ltd.) to remove high molecular weight conjugates of IAA. 6M NaOH was added to a final concentration of 1 M and the sample hydrolysed for 1h at room temperature (Bandurski and Schulze 1977). Following hydrolysis the pH was adjusted to 3– 3.5 by adding glacial acetic acid and samples kept on ice, prior to SPE clean-up. Strata C18-E (55μm, 70A) columns were prewashed sequentially with 2 mL hexane, 2 mL methanol, 5 mL deionized water and 1 mL 1% acetic acid. Samples were loaded onto columns and washed 4 times with 1 mL deionized water. IAA was eluted with 1 mL ACN and transferred to vials and stored at −20°C. The remaining half samples were prepared for free IAA analysis as described by Kabir et al. (2020). The analysis of ^12^C:^13^C IAA was conducted using a triple quadrupole Liquid Chromatograph Mass Spectrometer (LCMS)-8050, (Shimadzu) with XBridge™ C18 3.5 μm, 2.1×50 mm column (Phenomenex). The chromatography solvent was 20% acetonitrile: 80% 0.01 M acetic acid at a flow rate of 0.2 mL/min. The nebulizing, heating and drying gas flow were 3 L/min, 10 L/min and 10 L/min, respectively. Interface temperature was 300°C, DL was 250°C and the heat block temperature was 400°C. The interface used a capillary voltage of 4 kV. The mass spectrometer was operated in multiple-reaction-monitoring mode (collision energy, 14.0 eV), transitions from *m/z* 174.10 to 130.10 for [^12^C_6_] and *m/z* 180.20 to 136.15 for [^13^C_6_] were monitored. A series of standard mixtures of [^13^C_6_] and unlabelled IAA in different ratios 10:1 to 1:10 were also assayed to confirm accuracy of quantitative analysis. Data were obtained from the average of two technical replicates first, then the average and the standard error of three biological replicates from each developmental stage.

## Results

### Both wheat and rice have many TGW6-like proteins

Our BlastP search of the wheat proteome (IWGSC RefSeq v1.1), using default parameters in EnsemblPlants, revealed 48 homologues with amino acid identities to TaTGW6 ranging from 43.5 to 98.8%. Their encoding genes are listed in Table 1. Most of the genes (34 out of 48) are found in clusters of tandem repeats in which some genes have truncated sequences, encoding proteins of approximately 200 amino acids compared to TaTGW6 which has 345 amino acids. The chromosomal location of five genes including *TaTGW6* is listed as unknown even though this gene was originally identified from a QTL on chromosome 4A. To investigate whether rice also has multiple TGW6-like proteins, a BlastP search of the rice proteome was then carried out. This discovered 11 homologues of OsTGW6 with 45.4-92.0% amino acid identity, encoded by genes listed in Table 2. Similar to the wheat *TGW6-*like genes, 10 are found in four tandem clusters and two are truncated sequences.

**Table 1.** List of TaTGW6 homologues including locus ID and location, size of gene product, amino acid identity to TaTGW6 and number of introns. Amino acid identity was compared using Emboss Needle

| Clade | Gene ID (IWGSC RefSeq v1.1) | Amino acids | Percent amino acid ID with TaTGW6 | Chromosome | Introns |
| --- | --- | --- | --- | --- | --- |
| I | TraesCSU02G223800/TaTGW6 | 345 | 100.0 | U | none |
|  | TraesCSU02G255900 | 189 | 97.4 | U | none |
|  | TraesCSU02G240600 | 189 | 96.8 | U | none |
|  | TraesCS1A02G026500 | 345 | 98.8 | 1A | none |
|  | TraesCS7D02G075900 | 345 | 98.0 | 7D | none |
|  | TraesCS7D02G076000 | 341 | 96.5 | 7D | none |
|  | TraesCS7D02G076500 | 190 | 95.3 | 7D | none |
|  | TraesCS7D02G076600 | 209 | 96.2 | 7D | none |
|  | TraesCS3A02G496900 | 350 | 85.0 | 3A | none |
|  | TraesCS3A02G496700 | 345 | 84.4 | 3A | none |
|  | TraesCS3B02G559100 | 345 | 83.9 | 3B | none |
|  | TraesCS3B02G558800 | 209 | 86.6 | 3B | none |
|  | TraesCS3D02G504000 | 345 | 85.3 | 3D | none |
|  | TraesCS3D02G503900 | 311 | 80.5 | 3D | none |
|  | TraesCS3B02G338500 | 343 | 74.2 | 3B | none |
|  | TraesCS3A02G298200 | 317 | 73.8 | 3A | none |
|  | TraesCS3D02G303900 | 320 | 73.4 | 3D | none |
|  | TraesCS2B02G281700 | 344 | 67.7 | 2B | none |
|  | TraesCSU02G104000 | 348 | 68.4 | U | none |
|  | TraesCS7A02G138800 | 305 | 59.4 | 7A | 1 |
|  | TraesCS7A02G138700 | 209 | 73.2 | 7A | none |
|  | TraesCS7A02G138300 | 209 | 74.2 | 7A | none |
|  | TraesCS7D02G139900 | 348 | 67.0 | 7D | none |
|  | TraesCS7D02G140000 | 348 | 68.4 | 7D | none |
|  | TraesCS7D02G140500 | 345 | 67.8 | 7D | none |
|  | TraesCS7D02G140600 | 307 | 57.5 | 7D | 1 |
|  | TraesCS7B02G040400 | 348 | 67.2 | 7B | 1 |
|  | TraesCS7B02G040900 | 348 | 67.5 | 7B | none |
| II | TraesCS3A02G256600 | 336 | 50.4 | 3A | 3 |
|  | TraesCS3B02G289800 | 336 | 50.4 | 3B | 2 |
|  | TraesCS3D02G256900 | 336 | 47.6 | 3D | 2 |
|  | TraesCS5B02G382800 | 341 | 48.1 | 5B | 2 |
|  | TraesCS5B02G383000 | 339 | 48.1 | 5B | 2 |
|  | TraesCS5D02G389100 | 355 | 46.0 | 5D | 2 |
|  | TraesCS2A02G078000 | 343 | 48.3 | 2A | 1 |
|  | TraesCS2B02G093000 | 343 | 48.0 | 2B | 1 |
|  | TraesCS2B02G092900 | 342 | 48.6 | 2B | 1 |
|  | TraesCS2D02G076000 | 343 | 48.0 | 2D | 1 |
|  | TraesCS2D02G076200 | 343 | 48.0 | 2D | 1 |
|  | TraesCS2D02G076300 | 346 | 46.8 | 2D | 1 |
|  | TraesCS2D02G076400 | 343 | 47.7 | 2D | 1 |
|  | TraesCSU02G189700 | 343 | 47.7 | U | 1 |
| III | TraesCS5A02G188200 | 363 | 46.1 | 5A | 2 |
|  | TraesCS5A02G188300 | 377 | 43.5 | 5A | 1 |
|  | TraesCS5B02G195300 | 346 | 48.3 | 5B | 2 |
|  | TraesCS5B02G195400 | 364 | 47.9 | 5B | 2 |
|  | TraesCS5D02G202700 | 346 | 48.2 | 5D | 2 |
|  | TraesCS5D02G202800 | 364 | 46.8 | 5D | 2 |
|  | TraesCS7D02G076300 | 361 | 44.3 | 7D | 2 |
| U-unknown |  |  |  |  |  |

**Table 2.** List of OsTGW6 homologues including locus ID and location, size of gene product, amino acid identity to OsTGW6 and number of introns. Amino acid identity was compared using Emboss Needle

| Clade | Gene ID | Amino acids | Percent amino acid ID with OsTGW6 | Chromosome | Introns |
| --- | --- | --- | --- | --- | --- |
| I | LOC_Os06g41850/OsTGW6 | 350 | 100.0 | 6 | none |
|  | LOC_Os06g41830 | 329 | 84.6 | 6 | none |
|  | LOC_Os06g41820 | 345 | 92.0 | 6 | none |
|  | LOC_Os07g35970 | 350 | 64.2 | 7 | none |
|  | LOC_Os07g36060 | 351 | 62.1 | 7 | none |
|  | LOC_Os07g36040 | 248 | 70.4 | 7 | none |
|  | LOC_Os07g35990 | 250 | 70.2 | 7 | none |
| II | LOC_Os01g50330 | 324 | 48.6 | 1 | 1 |
|  | LOC_Os08g34330 | 350 | 47.1 | 8 | 2 |
|  | LOC_Os08g07810 | 356 | 47.3 | 8 | 2 |
| III | LOC_Os09g20810 | 362 | 45.4 | 9 | 2 |
|  | LOC_Os09g20684 | 364 | 45.5 | 9 | 2 |

The protein phylogenetic tree in Fig. 1 shows the relationships between wheat and rice TGW6 homologues. This divides into three major clades; the largest clade (clade I) contains 27 wheat and six rice proteins with greater than 57% amino acid identity to TaTGW6 and OsTGW6, respectively. Within clade I, wheat and rice proteins form separate branches. The largest branch contains TaTGW6 plus 13 wheat homologues. Five of these have truncated sequences, but there appear to be eight full-length proteins with over 80% amino acid identity to TaTGW6. At least 10 proteins are encoded by genes located in clusters of tandem repeats. An additional small branch in clade I has three proteins with approximately 74% amino acid identity to TaTGW6, all encoded on chromosome 3. The final wheat branch in clade I has 11 proteins with 57.5-74.2% amino acid identity to TaTGW6. The rice proteins are divided into two branches, each encoded by a cluster of tandemly repeated genes. The cluster on chromosome 6 includes *OsTGW6* as well as two additional genes encoding proteins with at least 84% amino acid identity to OsTGW6. We also noted that all rice genes encoding proteins in clade I and also most wheat genes in this clade had no introns (Table 1 and 2).

**Fig. 1.**
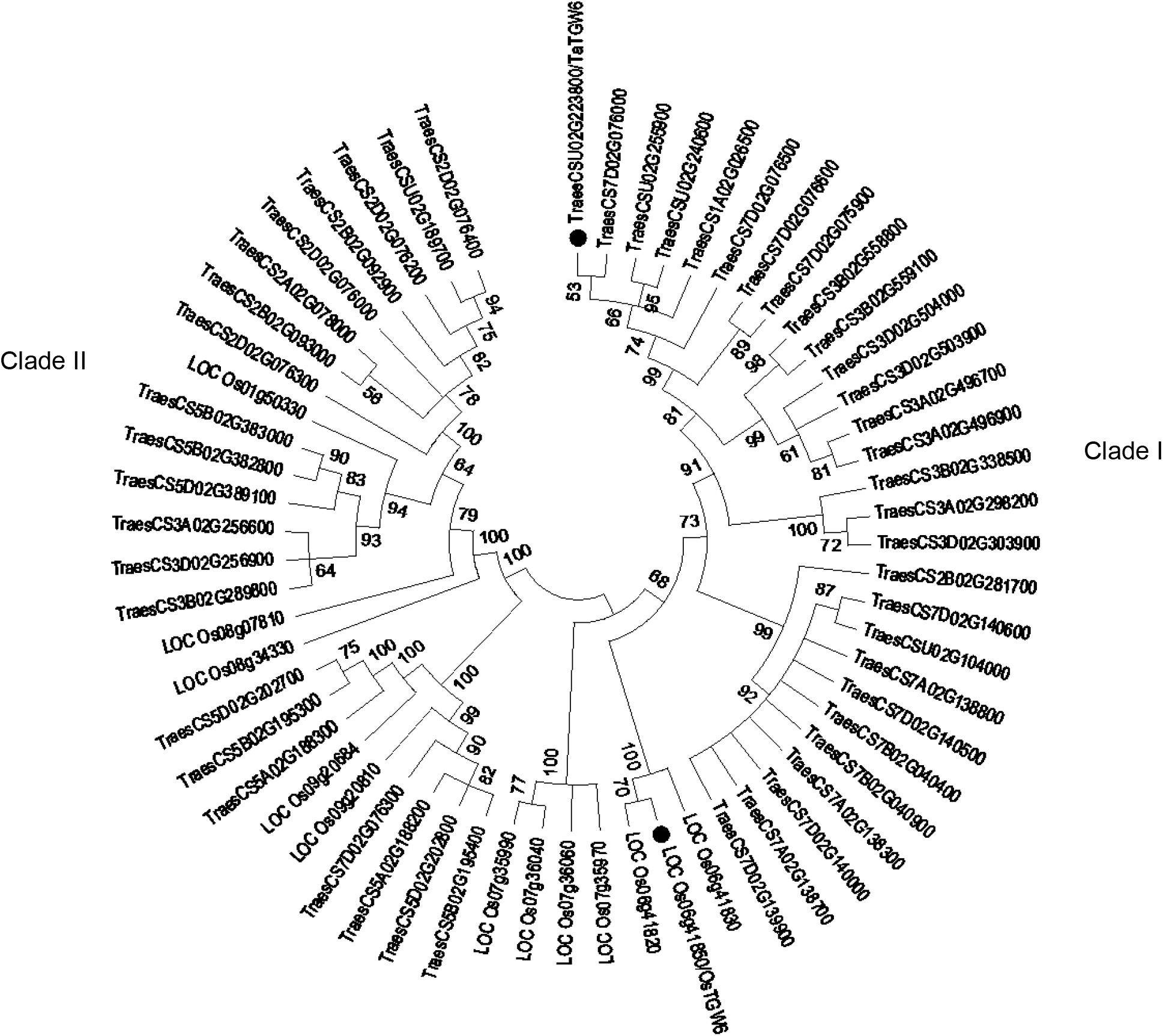
Phylogenetic tree showing relationships between TGW6 proteins from *Triticum aestivum* (Traes) and *Oryza sativa* (Os). The tree was constructed in MEGA7.0.26 (Kumar et al. 2016) using Maximum Likelihood method (Jones et al. 1992). Multiple sequence alignment was performed by MUSCLE (Edgar 2004). Bootstrap confidence levels were obtained using 500 replicates (Felsenstein 1985). Evolutionary distances were computed using Poisson correction method (Zuckerkandl and Pauling 1965). Black dots indicate TaTGW6 and OsTGW6.

Clade II contains 14 wheat and three rice proteins with 46-50% amino acid identity to TaTGW6 and OsTGW6. Clade III has seven full length wheat and two rice proteins. These have between 43-48% amino acid identity to TaTGW6 and OsTGW6. The genes encoding proteins in clades II and III are also primarily found in groups of tandem repeats in both wheat and rice.

### *TaTGW6, OsTGW6* and their clade I homologues are expressed in early inflorescence not in developing grains

Following reports that *TaTGW6* has maximum expression at 20 DAA whereas *OsTGW6* has highest expression at 2 DAA, we examined the expression of *TaTGW6* and 19 of its close homologues in clade I during grain development up to 20 DAA by reverse transcriptase PCR. Due to the very high homology within this large group of genes, primers were designed to amplify groups of similar genes as shown in Supplementary Table S1. Three separate primer sets including those from Hu et al. (2016a) were used for *TaTGW6* and its three closest homologues. Nevertheless no amplification was found from RNA samples derived from grain samples of any age. Poor RNA quality as well as non-functional primer sets were both ruled out as causes of the negative results via successful amplification of other IAA biosynthesis genes (*TAR* and *YUCCA*) and successful amplification from DNA template.

To further investigate the unexpected lack of expression in developing grains of wheat, we first investigated microarray data from PLEXdb (Dash et al. 2011) then RNA-seq data via expVIP for more information on gene expression. The RNA-seq database in particular has samples from many different tissues and experiments. Fig. 2 summarises information on the expression of *TaTGW6* and the 48 homologues listed in Table 1, throughout the plant from Chinese Spring samples, with genes organized by clade. These data confirm neither *TaTGW6* nor any of its clade I homologues have any expression in grains or in leaves or roots. Instead, *TaTGW6* and several *TaTGW6-*like genes show highly restricted expression only in the early inflorescence. In this tissue, highest expression was found for *TaTGW6*, TraesCS7D02G139900, TraesCS7D02G140500 and TraesCS7B02G040900. Most genes in clade II also have the same expression profile restricted to early inflorescence development. On the other hand, genes in clade III have three different expression patterns: TraesCS5B02G195400 and TraesCS5D02G202800 are expressed in a wider range of spike samples including anthers; TraesCS5B02G195300 and TraesCS5D02G202700 are upregulated in leaf tissue and young grains (2 DAA); TraesCS7D02G076300 is expressed in the stem and flag leaf. Only a single gene from clade II, TraesCSU02G189700, had any detectable expression in grains at 20 DAA and the highest expression of this gene was in the early inflorescence samples as for other clade II genes. We confirmed the expression of these clade II and III genes in nine-day-old seedling leaf, stamen and pistil, and 20 DAA in grains by RT-PCR. Additionally, our experimental data showed that mRNA for clade III genes, TraesCS5B02G195300 and TraesCS5D02G202700 is present in grains up to 5 DAA. However, we did not detect amplification of *TaTGW6* or its close homologues in RNA from these tissues.

**Fig. 2.**
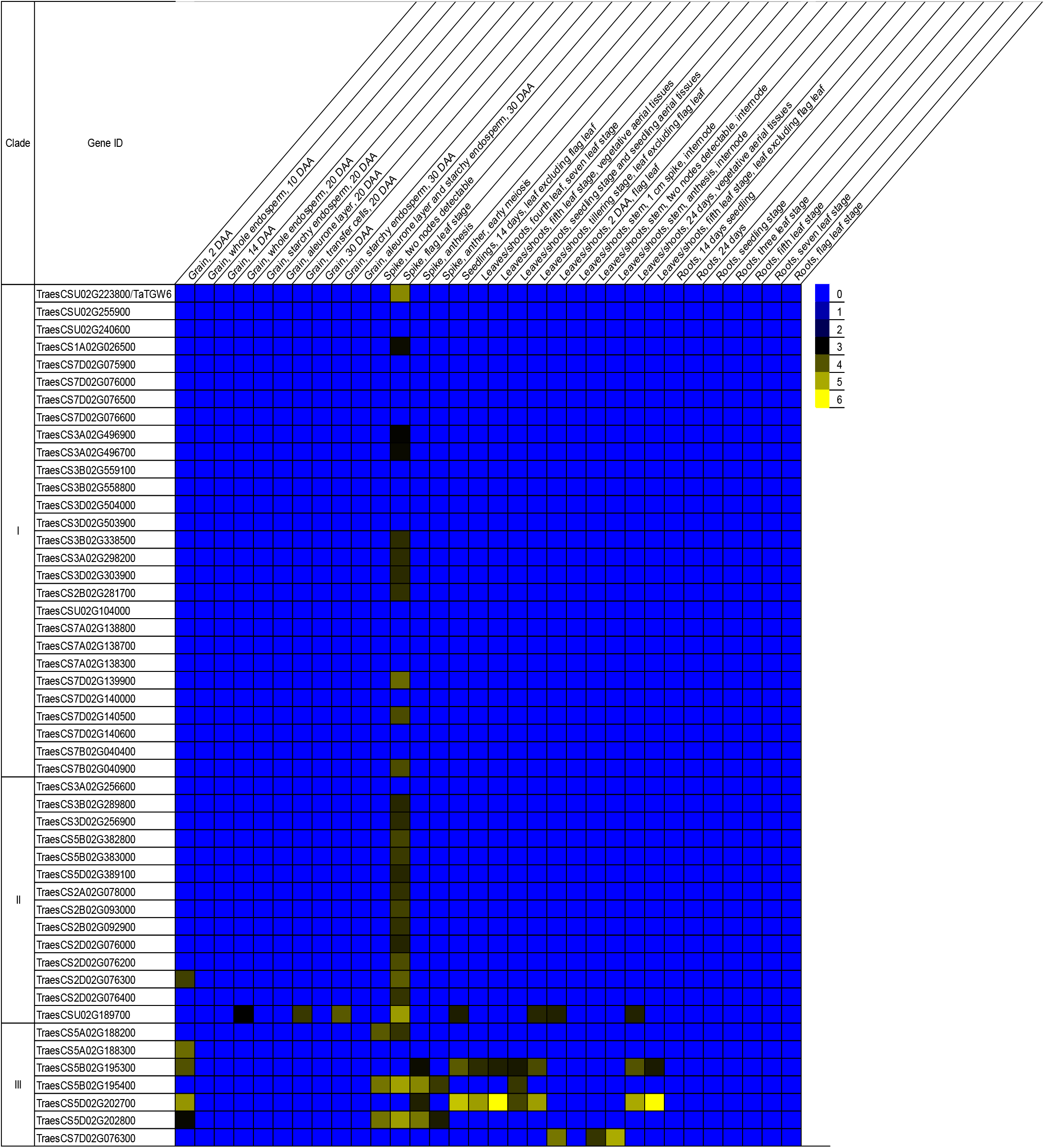
Heat map depicting expression of *TaTGW6* gene and its homologues based on RNA-seq data from Chinese Spring in expVIP (http://www.wheat-expression.com). The relative expression values are normalized in tpm (transcripts per million).

As these results differ radically from the published data, we also investigated the expression of *OsTGW6* and its homologues from rice, using microarray and RNA-seq databases. The RNA-seq results are presented in Fig. 3; both data sets confirmed that like *TaTGW6*, the rice gene, as well as its closest homologues on chromosome 6, is expressed only in the pre-emergent inflorescence. No expression was detected in grain samples. Other clade I genes on chromosome 7 are expressed both in the pre-emergent inflorescence as well as in anthers. The clade II genes on chromosome 8 are also expressed in pre-emergent inflorescence similar to their wheat orthologues. Only one gene in clade III, LOC_Os09g20684 is expressed in multiple tissues, particularly shoots. This gene appears to be orthologous to TraesCS5B02G195300 and TraesCS5D02G202700 (see Fig. 1) which have a similar expression profile.

**Fig. 3.**
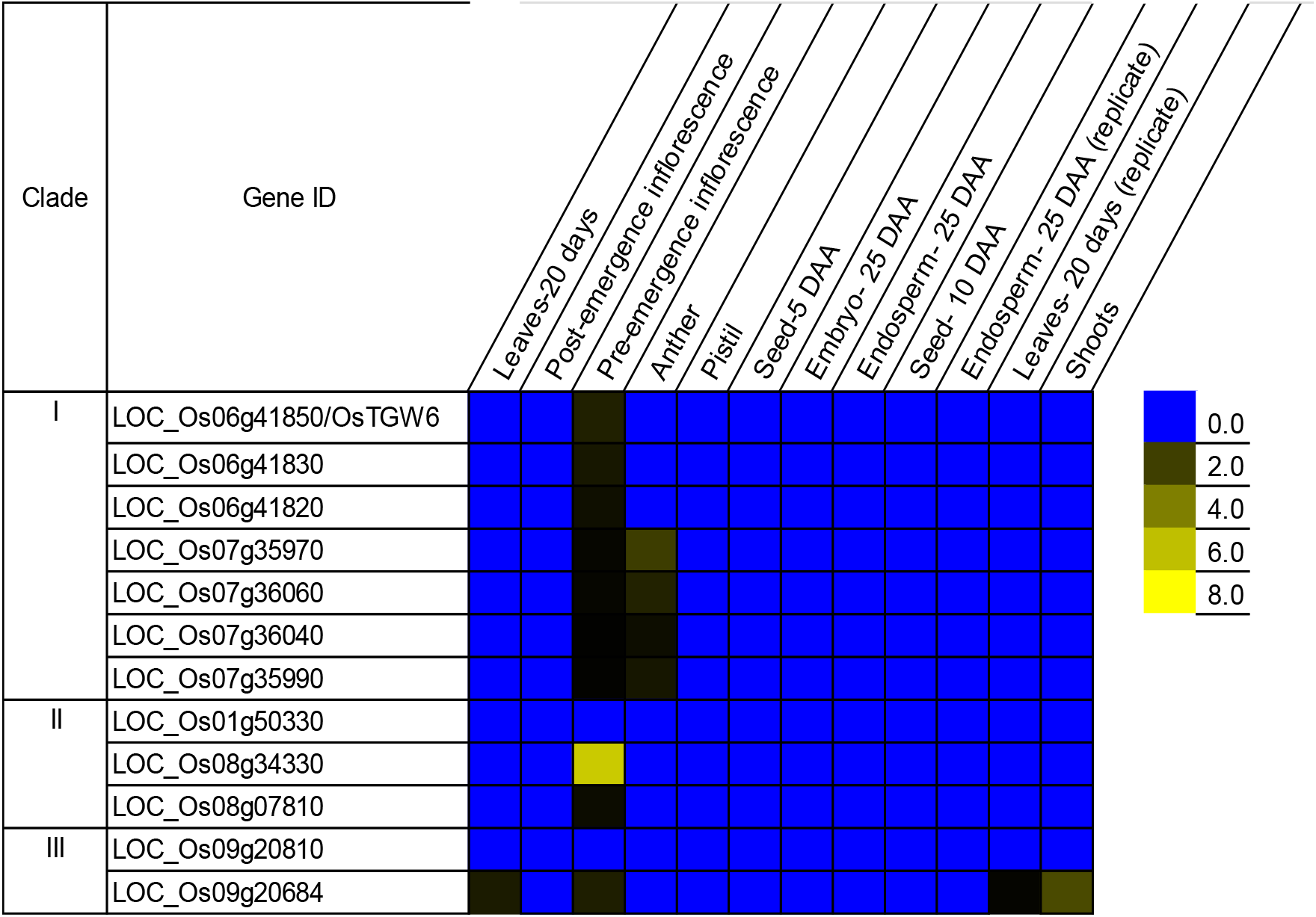
Heat map depicting expression of *OsTGW6* and its homologues based on RNA-seq data available in the Rice Genome Annotation Project (http://rice.plantbiology.msu.edu). Values are expressed as FPKM (Fragments Per Kilobase of transcript per Million mapped reads).

### Expression of *TaTGW6*-like genes varies between varieties and is similar to that of *OsSTRL2*

We noted a report in the literature that a more distant homologue of *TGW6* in rice, strictosidine synthase-like (*OsSTRL2*) (Zou et al. 2017), is also expressed in the pre-emergent inflorescence, specifically in the tapetum and microspores, and is required for normal pollen development. The expVIP data enabled us to compare expression of *TaTGW6-*like genes as well as the wheat orthologues of *OsSTRL2* (encoded by TraesCS4B02G215300, TraesCS4D02G215800, and TraesCS4A02G089000, Table 3) in Chinese Spring, other wheat varieties and in more specific microspore samples. Fig. 4 shows that *TaTGW6*, eight of its close homologues as well as the wheat orthologues of *OsSTRL2* are all expressed in microspores at the late vacuolated uninucleate stage, prior to mitosis. We noted that, *TaTGW6* and two other genes are more highly expressed than the putative *TaSTRL* genes. In addition, expression of *TaTGW6*-like genes varies between the four varieties for which data are available. *TaTGW6* itself appears not to be expressed in the variety Azhurnaya. However, the close homologue TraesCS3B02G559100 is expressed in these plants. The description of developmental stages varies between studies. However, *TaTGW6-*like gene activity appears to be restricted to full boot/spike “flag leaf stage” and is downregulated as the spike emerges from the boot. Expression of the *TaSTRL* genes is highest in the same samples, but continues a little later during spike emergence. However, *TaSTRL* genes are completely downregulated by anthesis.

**Table 3.** Putative wheat orthologues of *OsSTRL2* showing percent amino acid identity to OsSTRL and TaTGW6, compared using Emboss Needle

| Gene ID | Percent amino acid identity to<br>OsSTRL2 | Percent amino acid identity to<br>TaTGW6 |
| --- | --- | --- |
| TraesCS4A02G089000 | 82.0 | 30.9 |
| TraesCS4B02G215300 | 81.0 | 31.6 |
| TraesCS4D02G215800 | 81.4 | 32.0 |

**Fig. 4.**
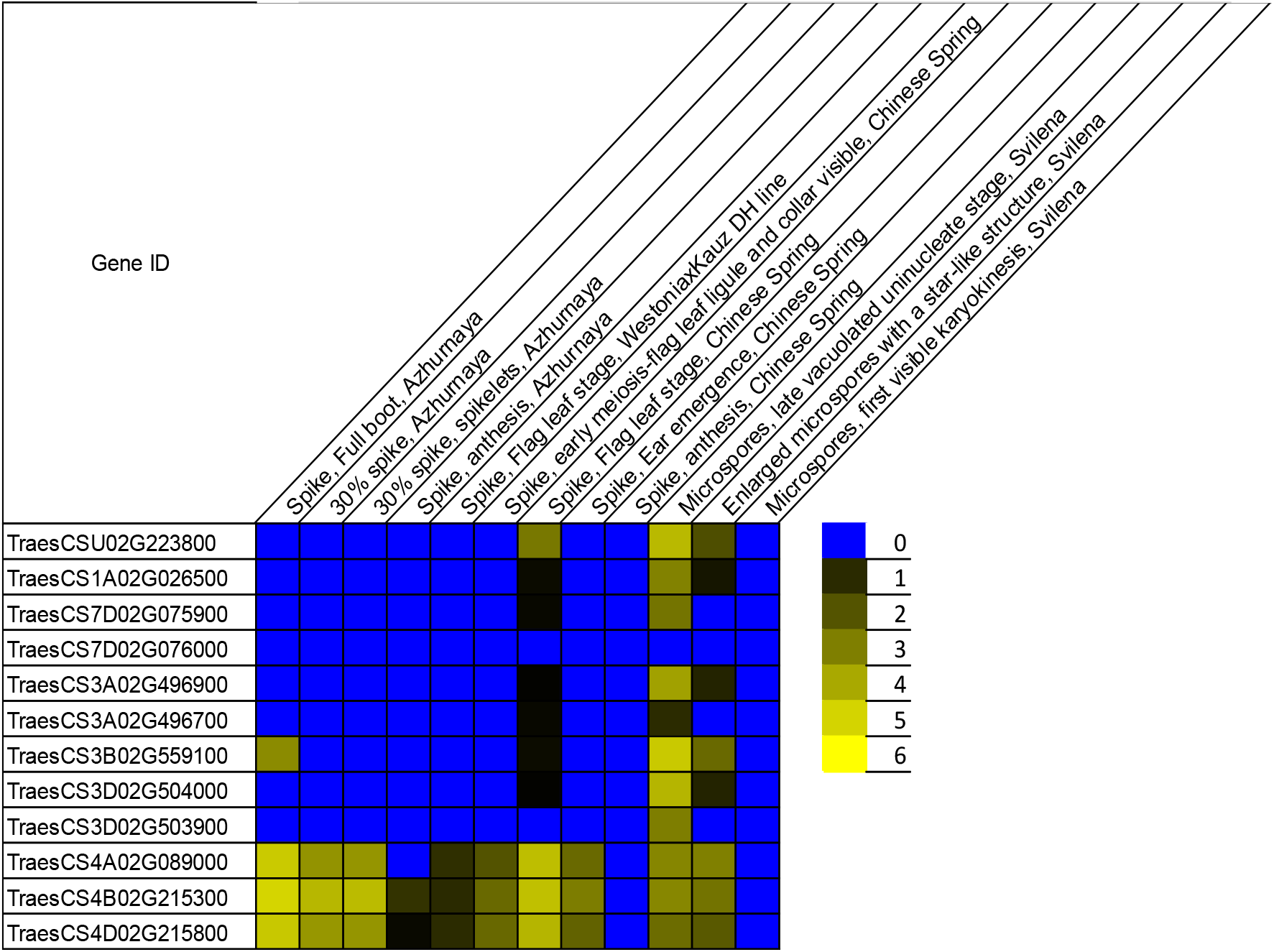
Heat map showing expression of full-length clade I *TaTGW6-*like genes as well as putative wheat orthologues of *OsSTRL2* in spike and microspore samples from four wheat varieties. The relative expression values are normalized in tpm (transcripts per million). Data come from three studies: Chinese Spring Development (Choulet et al. 2014), Azhurnaya development and spike drought (Ramírez-González et al. 2018), Microspores (Seifert et al. 2016) accessed via expVIP (http://www.wheat-expression.com).

### Investigation of existence and production of IAA-Glc in wheat grains

In the second part of the study, we investigated evidence for IAA-Glc (the proposed substrate of *TGW6*) as well as expression of genes (*IAGLU*) required for its production from IAA. A BlastP search of the wheat proteome with rice and maize IAGLU revealed a total of 38 homologous proteins. These form a large diverse family of UDP-glycosyltransferases (UGT) most of which have less than 50% amino acid identity to the query sequences. However, five proteins have 58.1-62.2% amino acid sequence identity to OsIAGLU and ZmIAGLU (Table 4) and are therefore putative TaIAGLUs. The phylogenetic tree of OsIAGLU, ZmIAGLU and their wheat homologues, rooted with the most similar homologue from rice, an uncharacterised putative UDP-glucosyl transferase (Fig. 5) indicates that the wheat sequences are likely to be conserved orthologues of OsIAGLU and ZmIAGLU.

**Table 4.** List of TaIAGLU homologues including locus ID, size of the gene product and amino acid identity to ZmIAGLU derived using Emboss Needle

| Gene ID<br>(IWGSC RefSeq v1.1) | Amino acids | Percent amino acid<br>ID with ZmIAGLU | Chromosome |
| --- | --- | --- | --- |
| TraesCS4D02G031500 | 500 | 58.1 | 4D |
| TraesCS4A02G279400 | 452 | 62.0 | 4A |
| TraesCS4D02G030900 | 453 | 62.1 | 4D |
| TraesCS3B02G282600 | 453 | 62.1 | 3B |
| TraesCS4B02G033000 | 504 | 59.0 | 4B |

**Fig. 5.**
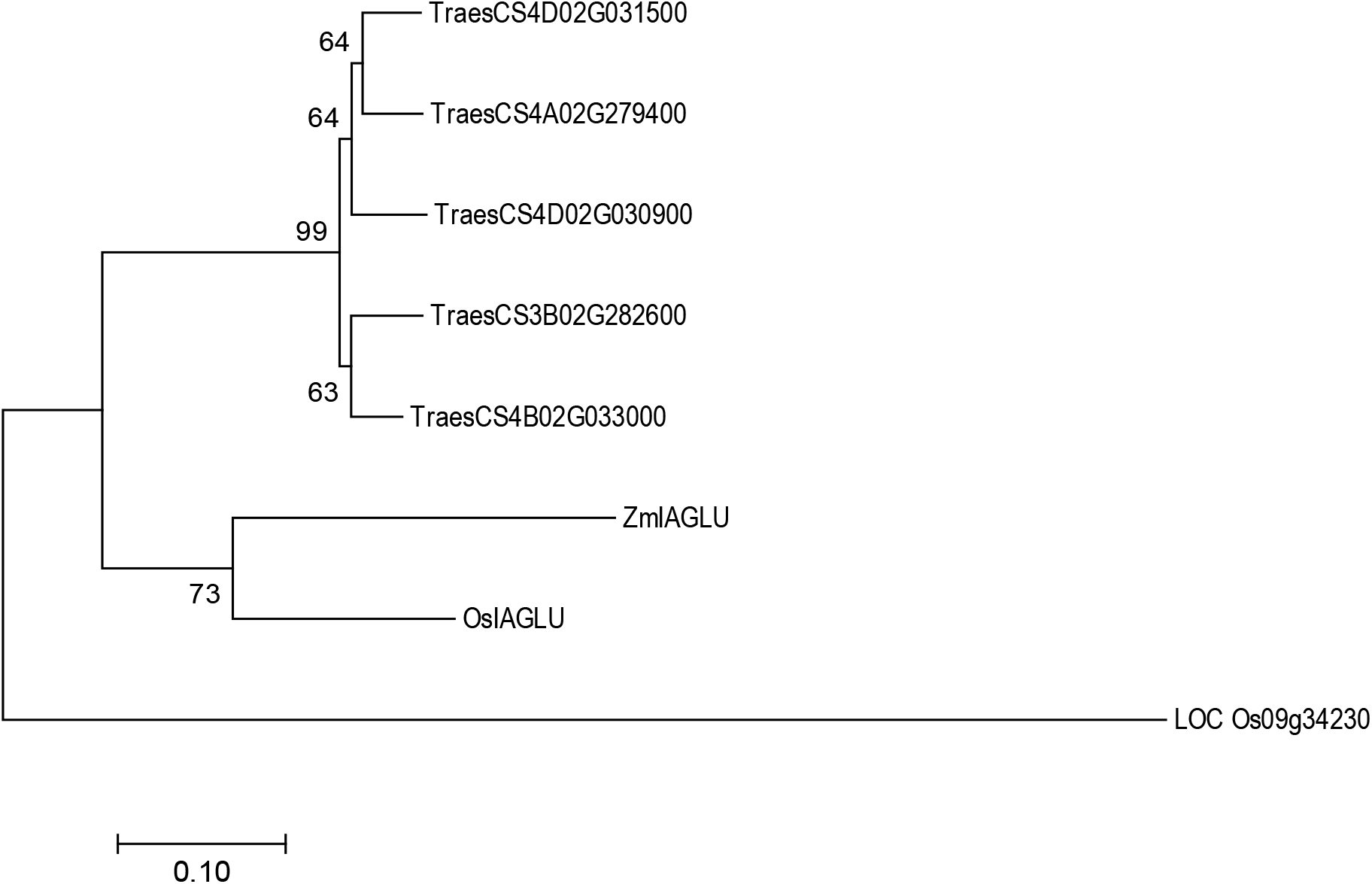
Phylogenetic tree showing relationships between IAGLU proteins from *Triticum aestivum* (Traes), *Oryza sativa* (Os) and *Zea mays* (Zm). The tree was constructed in MEGA7.0.26 (Kumar et al. 2016) using the Maximum Likelihood method (Jones et al. 1992). Multiple sequence alignment was performed by MUSCLE (Edgar 2004). Bootstrap confidence levels were obtained using 500 replicates (Felsenstein 1985). Evolutionary distances were computed using Poisson correction method (Zuckerkandl and Pauling 1965). The tree was rooted using LOC_Os09g34230 encoding a putative UDP-glucosyl transferase. Scale bar=0.1, amino acid substitutions per site.

*OsIAGLU* and *ZmIAGLU* expression has been reported in seedlings and in endosperm at 18 DAA, respectively. Thus we investigated *TaIAGLU* expression during grain development and in young seedlings by RT-PCR. However, we failed to detect any transcript for *TaIAGLU* genes in any sample. These results in combination with RNA-seq data (not shown), indicate that *TaIAGLU* is not expressed in developing grains or in the inflorescence.

To confirm the lack of availability of IAA-Glc as substrate for any hydrolase, we extracted and analysed low molecular weight ester IAA in young developing spikes as well as in grains at 20 and 30 DAA. Free IAA was found in all samples; pre-anthesis spikes (0.14 μg/g FW), 20 DAA (1.3 μg/g FW) and 30 DAA (0.57 μg/g FW). However, the amounts of IAA in samples that had been subjected to hydrolyse were the same as the free IAA values. Thus the bulk of IAA in all samples was free IAA, with very little low molecular weight ester IAA such as IAA-Glc.

Finally we also investigated whether wheat has any potential orthologues of UGT84B1, an UDP-glucose:indole-3-acetic acid glucosyltransferase from Arabidopsis encoded by AT2G23260. The BlastP search indicated that glycosyl transferases in wheat, as well as in rice and maize, have a maximum of 34% amino acid homology to UGT84B1 (data not shown) indicating that cereals do not have a conserved orthologue of this protein.

## Discussion

### *TaTGW6* and *OsTGW6* are part of a large gene family

In this study, we set out to re-evaluate the hypothesis that *TaTGW6* and *OsTGW6* affect grain size via their effects on IAA production in developing grains. In light of the hexaploid nature of wheat, we considered it surprising that single non-functional allele encoding an IAA-Glc hydrolase could have a measurable effect on IAA content and grain size. In addition, Hu et al. (2016a) had previously reported 95 sequences with high similarity to *TaTGW6* in a combination of wheat progenitor species and barley. We therefore investigated how many *TaTGW6-*like genes are present in the wheat genome and also expressed in developing wheat grains.

The BlastP search and subsequent phylogenetic analysis identified 48 homologues of *TaTGW6*, with most located in clusters of tandem repeats on particular chromosomes. A similar situation occurs in rice, in which *OsTGW6* and six close homologues are located in two clusters on chromosomes 6 and 7. The clusters of tandem repeats were most prevalent in clade I. In addition, most clade I genes had no introns. Finally we noted that although each clade contained proteins from both wheat and rice, branches in clades I and II contained proteins from a single species. All observations suggest that *TGW6-*like genes have undergone rapid and recent gene expansion.

This lineage-specific gene expansion is unexpected in a gene family involved in hormone regulation. Additionally, the existence of eight genes encoding proteins with more than 80% amino acid homology to TaTGW6 (two have above 95% identity), suggests that a single inactive allele is not likely to have a major effect on IAA content, unless only *TaTGW6* is expressed in grains. Similarly, rice has two genes on chromosome 6, encoding proteins with more than 84% amino acid homology to OsTGW6, suggesting that like wheat, a single non-functional allele in rice would be unlikely to have a significant impact on the IAA content of grains unless only *OsTGW6* is expressed.

### Expression data cast doubt on role of *TGW6* in grains

Although Hu et al. (2016a) found highest expression of *TaTGW6* at 20 DAA in grains and reported no effect of the inactive allele on IAA content of younger grains, the rice gene is reported to exert its effect on IAA levels at only 2 DAA. In addition, we have argued above that a single inactive allele could only reduce grain IAA content if its close homologues are not expressed in grains. Therefore, it was essential to confirm which *TGW6-*like genes are expressed and in what tissues and stage of development. To our surprise, we were not able to amplify transcripts of any genes in clade I, even with the use of primers from the Hu et al. (2016a) publication. Careful checking of these results included confirming the efficacy of primers with DNA as well as our ability to amplify clade III transcripts from the same RNA samples. The lack of expression in grains at any age was then confirmed using microarray and RNA-seq data as this became available. The RNA-seq data demonstrated that expression of *TaTGW6* is restricted to pre-emergent spike tissue. Close homologues of *TaTGW6* were also expressed in the same tissues. Subsequent examination of transcript information for rice genes confirmed the same situation. *OsTGW6* is not expressed in grain samples; instead, *OsTGW6* as well as all clade I homologues are also expressed in the pre-emergent inflorescence. Thus our results show that published data from Ishimaru et al. (2013) and Hu et al. (2016a) on expression of both *OsTGW6* and *TaTGW6* appear to be incorrect and that restricted expression of *TGW6* in the pre-emergent inflorescence is conserved across the two cereal species. We note that as neither *TaTGW6* nor *OsTGW6* have introns, extreme care needs to be taken to avoid any DNA contamination of RNA samples used for RT-PCR and to include no reverse transcriptase controls in all work. Our findings indicate that *TGW6* in neither species can play a direct role in the production of IAA in developing grains and that some other explanation should be sought to explain any effect on grain size.

A more detailed examination of expression of *TaTGW6* and its close homologues in different wheat varieties confirmed the restricted expression to pre-emergent inflorescence and also suggested that expression occurs in the microspores prior to mitosis. Whether expression is specifically restricted to microspores requires further investigation. However, the extensive species-specific gene expansion, expression restricted to a particular stage of pollen development as well as variable expression of different *TGW6-*like genes between wheat varieties all point to a possible role of this gene group in pollen development or possibly reproductive compatibility. We then came across a publication on a more distant homologue of *OsTGW6*, *strictosidine synthase-like, OsSTRL2*, (Zou et al. 2017). Although OsSTRL2 has only 34.6% amino acid identity with OsTGW6, it is expressed at the same stage of development. In addition, Zou et al. (2017) showed that expression was restricted to the tapetum and microspores; furthermore knockout of *OsSTRL2* resulted in male sterility. The mutation had no observable effect until stage 9 of pollen development (early microspore development), after which the microspores became wrinkled and shrunken indicating a key role for *OsSTRL2* at this very specific stage of pollen development. Zou et al. (2017) also listed all members of the *strictosidine synthase-like* family, including *OsTGW6* which they designated as *OsSTRL7*. No connection was made to the *TGW6* gene and the work by Ishimaru et al. (2013). They did however note, from mined RNA-seq data, that *OsSTRL5, OsSTRL6* and *OsSTRL7* (LOC_Os06g41820, LOC_Os06g41830 and LOC_Os06g41850) showed a similar expression pattern to *OsSTRL2*, though the signal was much weaker. In contrast, we noted that in the more specific microspore samples represented in the wheat RNA-seq database, *TaTGW6* was more highly expressed than *TaSTRL* genes. The *TaSTRL* genes were however expressed for a longer duration than *TaTGW6*. In summary, these data provide circumstantial evidence that *TGW6* in wheat and rice may play a similar role in pollen development to *OsSTRL2*.

### IAA-Glc is not present or produced in developing wheat grains

The second question addressed by this study was: is IAA-Glc present in developing wheat grains? TGW6 was reported to affect the auxin content of grains via IAA-Glc hydrolase activity. However, the existence of IAA-Glc was not investigated by either Ishimaru et al. (2013) or Hu et al. (2016a). Nevertheless mature seeds of several cereals including wheat are known to store IAA as ester conjugates with carbohydrates (Bandurski and Schulze 1977); these are hydrolysed to serve as a source of IAA during germination (Epstein et al. 1980). Ester conjugates of IAA have been extensively studied in maize, where the bulk are high molecular weight esters or esters of *myo*-inositol (Cohen and Bandurski 1982). Immature maize kernels are a rich source of IAA:UDP-glucose transferase enzyme activity; although, the resulting IAA-Glc is then transacetylated to *myo*-inositol (Cohen and Bandurski 1982). In addition, the *IAGLU* genes encoding this enzyme have been characterised from maize and rice, (Choi et al. 2012; Szerszen et al. 1994). As a prelude to analysis of IAA-Glc from developing wheat grains, we first assayed low molecular weight IAA ester conjugates via a well-established hydrolysis method. The comparison between free IAA and hydrolysed low molecular weight ester IAA indicated that the amount of the latter was negligible in comparison to free IAA. In addition, an exhaustive search for expression of the wheat orthologues of *IAGLU* was unable to detect gene activity in developing grains and other tissues. In contrast, our recent work on the TAR/YUCCA pathway showed this was highly active in developing wheat grains from 10 to 20 DAA (Kabir et al. 2020) similar to previously published work on rice (Abu-Zaitoon et al. 2012). Expression of *TaTAR2.3-1B, TaYUC9-1* and *TaYUC10* was strongly correlated with the free IAA content of developing wheat grains. We therefore conclude that *de novo* synthesis from tryptophan rather than hydrolysis of IAA-Glc is the major source of IAA during wheat grain development.

## Conclusion

Although *TaTGW6* and *OsTGW6* are widely cited to affect grain size in wheat and rice via their effects on IAA-Glc hydrolysis, we have demonstrated that neither gene is expressed in developing grains. In addition, the proposed substrate, IAA-Glc is not present in grains and the gene for its formation is not expressed in grains or spike tissue. We conclude that like maize, the IAA content of developing wheat grains is controlled by the expression of *TAR* and *YUCCA* genes. *TaTGW6* and *OsTGW6* are part of a large gene family that has undergone recent gene expansion, with many close homologues present in both species. *TGW6* and *TGW6-*like genes are expressed exclusively in the pre-emergent inflorescence of both wheat and rice. In wheat, expression occurs in the microspores prior to mitosis. Gene expression is closely similar to the homologous *OsSTRL2* and its wheat orthologues. We therefore suggest that *TaTGW6* and *OsTGW6* are most likely to play a role in pollen development.

## Declarations

### Availability of data and material

Not applicable

### Code availability

Not applicable

### Key message

Phylogenetic and expression analyses of grain weight genes *TaTGW6* and *OsTGW6*, and investigation of substrate availability indicate *TGW6* does not regulate auxin content of grains but may affect pollen development.

## Acknowledgements

The authors are grateful to Kirsten Drew for the analysis of IAA. The authors also acknowledge the Australian Government for providing a Research Training Program (RTP) PhD scholarship to Muhammed Rezwan Kabir.

## Author contribution statement

MRK and HMN conceived and designed the research. MRK performed all the experiments. MRK and HMN wrote the manuscript. Both authors read and approved the manuscript.

