## Supplementary data for "Reinvestigation of grain weight genes *TaTGW6* and *OsTGW6* casts doubt on their role in auxin regulation in developing grains"

**Supplementary Table S1** Primer sequences used for expression analysis of *TaTGW6* and *TaIAGLU* homologues in wheat tissues

| Gene ID | Primers | Product size (bp) |
|---|---|---|
| *TaTGW6*-like genes | | |
| TraesCSU02G223800/*TaTGW6*<br>TraesCS7D02G076000<br>TraesCS7D02G076600<br>TraesCS7D02G075900 | F: 5'-CCGATGCGTACAAAGGGCTT-3'<br>R: 5'-TGACCGGTGACTTGATCGAC-3' | 127 |
| TraesCSU02G223800/*TaTGW6*<br>TraesCS7D02G076000<br>TraesCS1A02G026500<br>TraesCS7D02G075900 | F: 5'-CGATGCGTACAAAGGGCTTA-3'<br>R: 5'-TTGACCGGTGACTTGATCGA-3' | 127 |
| TraesCSU02G223800/*TaTGW6*<br>TraesCSU02G255900<br>TraesCSU02G240600<br>TraesCS1A02G026500<br>TraesCS7D02G075900 | *F: 5'-CACCTCGTGGTCGCATCT-3'<br>*R: 5'-ATCTGGGTAGCCCGGCAG-3' | 108 |
| TraesCS3A02G496900<br>TraesCS3A02G496700<br>TraesCS3D02G504000<br>TraesCS3B02G559100 | F: 5'-CCACTAGCTTCGATGCCTCA-3'<br>R: 5'-CCGGGTCCATATGCATAGGT-3' | 181 |
| TraesCS3B02G338500 | F: 5'-CAGGTCACCGGTCAAGTCTA-3'<br>R: 5'-CAATCAGATGCGTACGGTCG-3' | 199 |
| TraesCS3A02G298200 | F: 5'-ACCGACCGTACGCATCTAAT-3'<br>R: 5'-CCCAGTAACCTCCCCTTCTG-3' | 154 |
| TraesCS2B02G281700 | F: 5'-CTAAAGGCCGGCATCACTTAC-3'<br>R: 5'-AGTACCTCAACAACTTGCACG-3' | 100 |
| TraesCS7D02G140600 | F: 5'-CGCGTGCTGAAGTGGAAC-3'<br>R: 5'-GCAGTCTCCGGGCGAA-3' | 107 |
| TraesCS7A02G138800 | F: 5'-GGGCACGATGGAGCTATTTG-3'<br>R: 5'-AACATCGATCCTCAGGGCAA | 151 |
| TraesCS7D02G140500 | F: 5'-CAATGTGAGGCCCGACAAAA-3'<br>R: 5'-TCGATCCTCAGTGCTAGCAG | 105 |
| TraesCS7B02G040400<br>TraesCS7B02G040900<br>TraesCS7D02G139900 | F: 5'-AAAGGCCAGGGTCCATACAG-3'<br>R: 5'-TGCTGTAGTCGGGGTTGTAG-3' | 103 |
| TraesCS5B02G195300<br>TraesCS5D02G202700<br>TraesCS5A02G188300 | F: 5'-GGCTGCAGTTCCACCA-3'<br>R: 5'-CACCACCAGTAGGTACTCGCT-3' | 224 |
| TraesCS5D02G202800<br>TraesCS5B02G195400<br>TraesCS5A02G188200 | F: 5'-TCTACCAACGGAGCGAGTACA-3'<br>R: 5'-CTGAGCACGGTGACGTT-3' | 100 |
| TraesCSU02G189700<br>TraesCS2D02G076400 | F: 5'-CCACGAAGGTTTAATACGGAAA-3'<br>R: 5'-GCCTTTGGTCCCTGGAGATA-3' | 209 |
| Putative *TaIAGLU* genes | | |
| TraesCS4A02G279400<br>TraesCS4D02G031500 | F: 5'-GTGGGCTGCTTCGTCAC-3'<br>R: 5'-GTTGATCGGCTGGTCGGT-3' | 105 |
